# Molecular responses to freshwater limitation in the mangrove tree *Avicennia germinans* (Acanthaceae)

**DOI:** 10.1101/731760

**Authors:** Mariana Vargas Cruz, Gustavo Maruyama Mori, Dong-Ha Oh, Maheshi Dassanayake, Maria Imaculada Zucchi, Rafael Silva Oliveira, Anete Pereira de Souza

## Abstract

Environmental variation along the geographical space can shape populations by natural selection. In the context of global warming, accompanied by substantial changes in precipitation regimes, it is crucial to understand the role of environmental heterogeneity in tropical trees adaptation, given their disproportional contribution to water and carbon biogeochemical cycles. Here we investigated how heterogeneity in freshwater availability along tropical wetlands has influenced molecular variations of the Black-Mangrove (*Avicennia germinans*). Fifty-seven trees were sampled in seven sites differing markedly on precipitation regime and riverine freshwater inputs. Using 2,297 genome-wide single nucleotide polymorphic markers, we found signatures of natural selection by the genotype association with the precipitation of the warmest quarter and the annual precipitation. We also found candidate loci for selection, based on statistical deviations from neutral expectations of interpopulation differentiation. Most candidate loci present within coding sequences were functionally associated with central aspects of drought-tolerance or plant response to drought. Complementarily, our results suggest the occurrence of rapid evolution of a population, likely in response to sudden and persistent limitation in plant access to soil water, following a road construction in 1974. Observations supporting rapid evolution included reduction in tree size and changes in allele frequencies and in transcripts expression levels associated with increased drought-tolerance, through accumulation of osmoprotectants and antioxidants, biosynthesis of plant cuticle, protection against stress-induced proteins degradation, stomatal closure, photorespiration and photosynthesis. We describe a major role of spatial heterogeneity in freshwater availability in the specialization of this typically tropical tree.

## 1. Introduction

Natural ecosystems are characterized by wide environmental heterogeneity over the geographical space. These spatial variations in several abiotic conditions can shape populations specializations, particularly in widespread species, through changes in frequencies of genotypes and phenotypes (Kawecki & Ebert, 2004). Environmental gradients are, therefore, natural laboratories for the study of environmental selection (De Frenne et al., 2013). As sessile organisms, plants are incapable of escaping from unfavorable conditions and, therefore, are ideal models for investigating adaptation through phenotypic plasticity and/or adaptive genetic variation. They are often subject to a wide range of environmental factors, such as water, light, temperature and nutrients availability. Among the various environmental factors that determine adaptive phenotypic and genotypic diversity in plants, freshwater availability has a prominent role (Choat et al., 2018; Phillips et al., 2010). Accordingly, population differentiation in drought-tolerance has been widely identified in various studies, providing valuable insights on evolutionary consequences of spatial variation in freshwater availability in plant species (Aranda et al., 2014; Donovan, Ludwig, Rosenthal, Rieseberg, & Dudley, 2009; Etterson, 2004; Heschel & Riginos, 2005; Keller et al., 2011; Ramírez-Valiente et al., 2018). Reduced freshwater availability resulting from the combination of high atmospheric temperature with low rainfall and air humidity represent a major threat to plants, especially trees and forest ecosystems they form (Asner et al., 2016; Bennett, McDowell, Allen, & Anderson-Teixeira, 2015; McDowell & Allen, 2015). These changes influence major components of resource-use: it reduces the soil water potential, which limits the water and nutrients supply to leaves, and raises the air vapor pressure deficit (VPD), increasing water loss through transpiration (McRae, 1980; Novick et al., 2016). In response to these conditions, plants close their stomata (McAdam & Brodribb, 2015; Tyree & Sperry, 1989), decreasing the chance of death from hydraulic failure (Rowland et al., 2015), despite causing negative impacts on photosynthesis and productivity (Lawlor, 2002).

Currently, there is a great interest in understanding genotypic and phenotypic basis of trees resistance to drought, as these mechanisms are key to improve predictions of environmental consequences of extreme events and to elaborate plans to mitigate forest loss (Corlett, 2016; da Costa et al., 2010; Phillips et al., 2009). Substantial advances in the understanding of phenotypic characteristics that enhance drought-tolerance in trees have been achieved recently (Bartlett, Scoffoni, & Sack, 2012; Bennett et al., 2015; Hacke, Sperry, Wheeler, & Castro, 2006; Powell et al., 2017), however little is known about the molecular basis of water-stress tolerance, particularly in tropical species (Holliday et al., 2017), which contribute disproportionally to global carbon cycle (Corlett, 2016).

In this study, we investigated the role of environmental selection along a gradient of freshwater availability in shaping the molecular variation of a typically tropical and abundant tree, *Avicennia germinans* (L.) L. (Acanthaceae). The species is particularly suitable for the study of mechanisms involved in trees adaptation to drought, since it is the most widespread mangrove in the Atlantic East-Pacific biogeographic region (Ellison, Farnsworth, & Merkt, 1999), belonging to the most tolerant mangrove genus to limiting conditions for water acquisition, as drought, freezing and salinity (Pranchai et al., 2017; Reef & Lovelock, 2015; Stuart, Choat, Martin, Holbrook, & Ball, 2007). These trees occur naturally under daily and seasonal fluctuations in temperature, air humidity, freshwater inputs and soil salinity (Tomlinson, 1986). In tropical arid zones or during dry seasons, VPD, soil salinity and water-deficits can reach extreme levels, hindering the maintenance of water and ion homeostasis and limiting carbon gain and plant growth (Clough, Sim, Inlet, Bay, & Rivers, 1989; Lin & Sternberg, 1992).

We hypothesized that spatial heterogeneity in freshwater availability shapes adaptive variation in allele frequencies and in expression profiles of transcripts associated with the response to and tolerance of limiting freshwater in tropical trees. We sampled *A. germinans* individuals along a wide longitudinal range in an equatorial region (from 0°43’12” S to 8°31’48” S of latitude), showing narrow spatial heterogeneity in temperature and solar radiation, but encompassing high variation in the intensity and duration of the dry season, in annual precipitation levels and in riverine freshwater input, which influence levels of soil salinity (Figure 1). We used the Nextera-tagmented reductively-amplified DNA (nextRAD) sequencing approach for the identification and genotyping of genome-wide single nucleotide polymorphisms (SNPs). We analyzed the organization of the genetic structure and performed statistical tests to identify candidate loci for selection. To minimize false positives in the detection of candidate loci (Lotterhos & Whitlock, 2015), we used distinct approaches: (1) based on the identification of loci deviating from neutral models of interpopulational genetic variation (F_ST_ outlier tests) and (2) based on direct genetic-environment (G-E) association tests. RNA sequencing (RNA-Seq) was used to assemble and characterize the transcriptome of the species, providing a functional basis for the annotation of candidate loci for selection. Additionally, we examined the role of molecular adaptation to extreme soil freshwater limitation via differential gene expression analysis between samples from adjacent sites differing in tidal inundation frequency (Lara & Cohen, 2006; Pranchai et al., 2017), acclimated in pots under homogeneous, watered conditions. Our results provide converging signs of adaptive responses of *A. germinans* to the environmental heterogeneity in freshwater availability, including a case suggesting the rapid evolution of a population, with changes in phenotype and in the genetic profile. We highlight the relevance of environmental heterogeneity in freshwater availability as a key selective pressure in tropical tree species.

**Figure 1.**
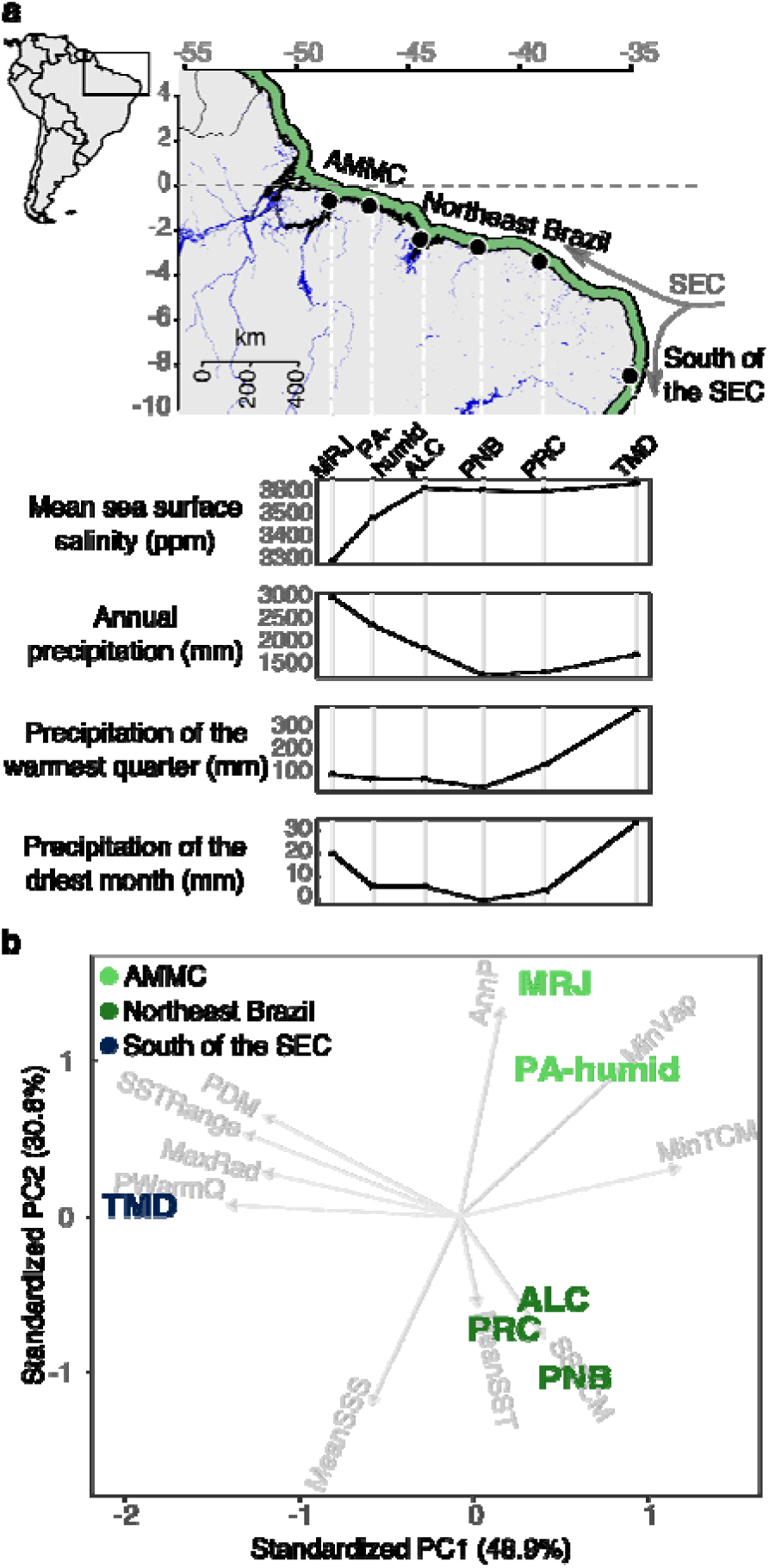
*Avicennia germinans* sampling sites along a low-latitude salinity and precipitation gradient. (a) *Top:* Geographical location of the study area and sampling sites. Black points represent sampling sites, green area represents the occurrence of the species, and blue areas represent ponds and rivers. *Bottom:* Environmental variation based on sea surface salinity and precipitation variables across sampling sites (source: WorldClim and Marspec). *AMMC*: Amazon Macrotidal Mangrove Coast. *SEC*: South Equatorial Current. (b) Bidimensional projection of sampling sites, based on a principal component analysis. PC1 and PC2 retained 79.7% of the variance of non-collinear environmental variables (eigenvectors). *AnnP*: Annual precipitation. *MinVap*: Minimum air water vapor pressure. *MinTCM*: Minimum temperature of the coldest month. *SSTCM*: Sea surface temperature of the coldest month. *MeanSST*: Mean sea surface temperature. *MeanSSS*: Mean sea surface salinity. *PWarmQ*: Precipitation of the warmest quarter. *MaxRad*: Maximum solar radiation. *SSTRange*: Sea surface temperature range. *PDM*: Precipitation of the driest month.

## 2. Materials and methods

### 2.1 Study system

*Avicennia germinans* (L.) L. is one of the most widespread mangroves in the Atlantic-East Pacific biogeographic region. It presents a generalist insect pollination system (Nadia, Menezes, & Machado, 2013), and is predominantly outcrossing, although it supports self-fertilization (Mori, Zucchi, & Souza, 2015; Nettel-Hernanz, S.Dodd, Ochoa-Zavala, Tovilla-Hernández, & Días-Gallegos, 2013). The species reproduces by producing cryptoviviparous, buoyant, and salt-tolerant propagules, allowing for long-distance sea dispersal (Mori, Zucchi, Sampaio, & Souza, 2015; Tomlinson, 1986).

### 2.2 Study area

Sampling sites were located over more than 1,800 km of the north-northeast Brazilian coastline, between 0.724° S and 8.526° S of latitude, along a spatial gradient in freshwater availability (Figure 1a, Table 1). We obtained environmental data for each sampling site, consisting of 21 bioclimatic and oceanographic variables from the public databases WorldClim (Hijmans, Cameron, Parra, Jones, & Jarvis, 2005) (15 precipitation, radiation and air vapor pressure layers) and Marspec (Sbrocco & Barber, 2013) (four sea surface salinity and temperature layers). To minimize the collinearity of variables, we used a correlation threshold of 0.8 (Supplemental Figure S1). We classified the study area into three distinguishable regions, based on a principal component analysis (Figure 1b): (1) the Amazon Macrotidal Mangrove Coast (AMMC), the world’s largest continuous mangrove belt (Nascimento Jr., Souza-Filho, Proisy, Lucas, & Rosenqvist, 2013), which has a mean annual precipitation above 2,000 mm yr^-1^ and is influenced by the mouth of the Amazon River; (2) mangroves of Northeast Brazil, which show limited forest development (Schaeffer-Novelli, Cintrón-Molero, Adaime, & de Camargo, 1990) and is characterized by the lack of riverine freshwater inputs and a mean annual precipitation below 2,000 mm yr^-1^, with pronounced and long dry seasons, and less than 30 mm of precipitation in the driest quarter; and (3) a region influenced by the southward-flowing branch of the South Equatorial coastal current (South of the SEC), characterized by reduced riverine freshwater inputs, mean annual precipitation below 2,000 mm yr^-1^, but less pronounced dry season, showing more than 100 mm of precipitation in the driest quarter.

**Table 1.** Characterization and geographic location of *Avicennia germinans* sampling sites.

| Sampling site ID | Climate classification† | Latitude; Longitude | Number of samples |  |
| --- | --- | --- | --- | --- |
|  |  |  | nextRAD | RNA-Seq |
| MRJ | Am | 00.72 S; 48.49 W | 8 | - |
| PA-arid‡ | Am | 00.90 S; 46.69 W | 9 | 3 |
| PA-humid | Am | 00.94 S; 46.72 W | 9 | 3 |
| ALC | Aw | 02.41 S; 44.41 W | 5 | - |
| PNB | Aw | 02.78 S; 41.82 W | 9 | - |
| PRC | Aw | 03.41 S; 39.06 W | 7 | - |
| TMD | As | 08.53 S; 35.01 W | 10 | - |

†Köppen-Geiger climate classification system (Alvares, Stape, Sentelhas, Gonçalves, & Sparovek, 2013). *Am*: Tropical monsoon climate; *Aw*: Tropical wet savanna climate; *As*: Tropical dry savanna climate.
‡Samples from a forest flooded exclusively during spring tides (Lara & Cohen, 2006).

In the AMMC, we visited two adjacent sites, which require a more detailed description, both located in the peninsula of Ajuruteua, state of Pará, between the Maiaú and Caeté estuaries (Figure 2a). The peninsula was covered by a preserved mangrove forest (Cohen & Lara, 2003), until it was divided by the construction of a road (Figure 2b), in 1974. The hydrology part of the forest was dramatically changed, no longer being influenced by the Caeté River. Instead, it started flooding exclusively during the highest spring tides of the Maiaú River. These changes resulted in the forest dieback and subsequent recolonization of the impacted area, mainly by *A. germinans*. Pore water salinity accumulated to extremely high levels (100 ppt at 50-cm depth), and the air surface temperature frequently exceeds 40 °C (Vogt et al., 2014). These environmental features contributed to the dwarfism of recolonizing individuals (Cohen & Lara, 2003; Pranchai et al., 2017), whose shrub architecture (up to 2.0 m in height) contrasts to the former tall morphology (up to 30 m in height), still observed on surrounding areas, where the hydrology remained unaltered (Menezes, Berger, & Mehlig, 2008) (Figure 2b). Throughout this work, we refer to the western side of the road as ‘PA-arid’ and to the eastern side as ‘PA-humid’.

**Figure 2.**
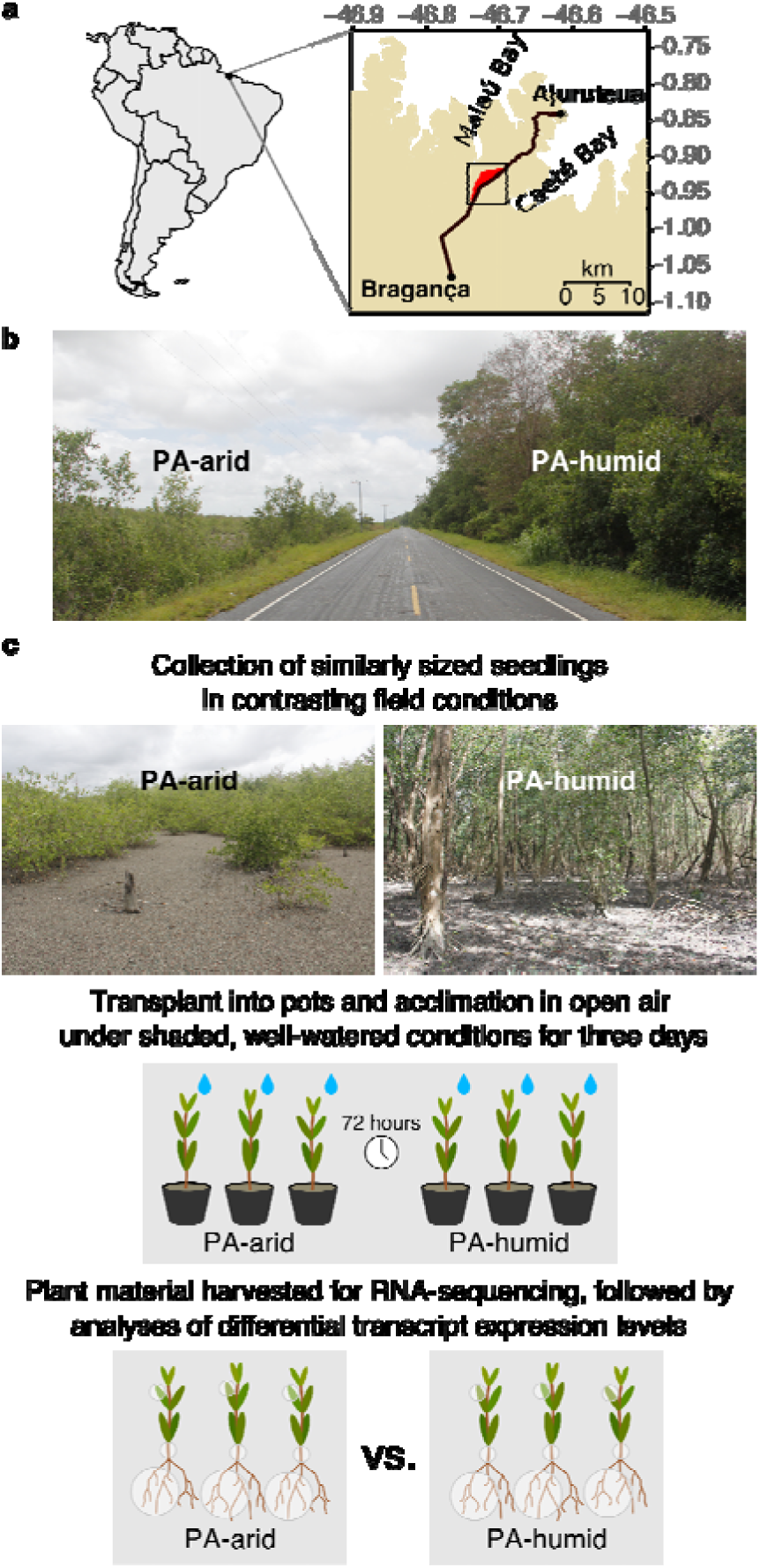
PA-arid and PA-humid sampling sites and RNA sequencing (RNA-Seq) experimental design. (a) Geographical location of sites, highlighted by a square (red colored area is the PA-arid site). (b) Photograph of a section of the Bragança-Ajuruteua road (Pará, Brazil) along which severe changes in hydrology altered the mangrove community and tree morphology. (c) RNA-Seq experimental design. *Photographs authors:* G.M. Mori and M.V. Cruz.

### 2.3 DNA extraction and sequencing

Leaves from 57 adult *A. germinans* trees were sampled in seven distinct sites (Table 1) and stored in bags with silica gel. DNA extraction was performed using the DNeasy Plant Mini Kit (QIAGEN) and NucleoSpin Plant II (Macherey Nagel). DNA quality and quantity were assessed using 1% agarose gel electrophoresis and QuantiFluor dsDNA System in a Quantus fluorometer (Promega). NextRAD libraries were constructed by SNPsaurus (SNPsaurus, LLC) (Russello, Waterhouse, Etter, & Johnson, 2015). Genomic DNA fragmentation and short-adapter ligation were performed with Nextera reagent (Illumina, Inc.), followed by amplification, in which one of the primers matched the adapter and extended by nine arbitrary nucleotides of DNA. Thus, amplicons were fixed at the selective end, and their lengths depended on the initial fragmentation, leading to consistent genotyping of amplified loci. Subsequently, nextRAD libraries were sequenced in a HiSeq 2500 (Illumina, Inc), with 100-bp single-end chemistry (Supplemental Figure S2).

### 2.4 SNP detection and genotyping

Genotyping-by-sequencing used custom scripts (SNPsaurus, LLC) to create a reference catalog of abundant reads. Read mapping to the catalog allowed two mismatches. Biallelic loci present in at least 10% of samples were selected for the following steps. Using VCFtools 0.1.12b (Danecek et al., 2011), we selected high-quality sequences (Phred score > 30), allowing a maximum of 65% missing data and one SNP per sequence, requiring a minimum coverage of 8x and a minor allele frequency ≥ 0.05 (Supplemental Figure S2). To reduce false SNP detection rates due to paralogy or low-quality genotype calls, we used a maximum read of 56, resulting from the product of the average read depth and 1.5 standard deviation of the mean.

### 2.5 Genetic diversity analysis and identification of candidate SNP loci associated with water-stress tolerance

For each sampling site, we estimated the genetic diversity using per-site nucleotide diversity (π), calculated using VCFtools (Danecek et al., 2011); private alleles (pA), observed (H_O_) and expected (H_E_) heterozygosities, using poppr 2.8.2 (Kamvar, Tabima, & Grünwald, 2014); and the percentage of polymorphic loci (%Poly), using adegenet 2.1.1 (Jombart & Ahmed, 2011). We also estimated pairwise genetic differentiation (F_ST_), the inbreeding coefficient (F_IS_) and its 95% confidence interval, through 1000 bootstrap resampling over all SNP loci, using hierfstat 0.04-22 (Goudet, 2005). The genetic structure of *A. germinans* was described by a multivariate model-free method, the discriminant analysis of principal components (DAPC) (Jombart, Devillard, & Balloux, 2010), and ADMIXTURE 1.3.0 (Alexander, Novembre, & Lange, 2009). We considered numbers of groups from 1 to 50 and the Bayesian information criteria to determine the number of groups (K). The optim.a.score function was used to avoid overfitting during discrimination steps. For ADMIXTURE analysis, we performed three separate runs for values of K varying from 1 to 15, using the block-relaxation method for point estimation. Computing was terminated when estimates increased by less than 0.0001. The lowest level of cross-validation error indicated the most likely K-value.

To detect signatures of natural selection, we used three distinct methods. Loci that were highly correlated with the environmental structure (false discovery rate (FDR) <0.05) were detected using the R package LEA (Frichot & François, 2015), which is based on analyses of genetic structure and ecological association (G-E association tests). In these analyses, we used non-collinear environmental bioclimatic and oceanographic variables (Supplemental Figure S1) retrieved from the public databases WorldClim (Hijmans et al., 2005) (precipitation, radiation and air vapor pressure layers) and Marspec (Sbrocco & Barber, 2013) (sea surface salinity and temperature layers). Since the environmental divergence between PA-arid and PA-humid is so evident (Figure 2), yet undistinguishable based on highest-resolution environmental datasets, we excluded PA-arid samples from G-E association tests.

To overcome limitations in the acquisition of environmental data for the PA-arid site, two other methods were used to detect candidate loci for selection. These methods are solely based on deviations from neutral expectations of allele frequency distributions, regardless of environmental variation data across sampling sites (F_ST_ outlier tests). With Pcadapt 2.0 (Luu, Bazin, & Blum, 2016), population structure was determined by a principal component analysis. A disproportional relation to this structure (FDR < 0.05) was indicative of selection. With Lositan (Antao, Lopes, Lopes, Beja-Pereira, & Luikart, 2008), which uses the FDIST2 method (Beaumont & Nichols, 1996), we assessed the relationship between F_ST_ and H_E_ to describe the neutral distribution under an island migration model. Hence, we detected loci with excessively high or low F_ST_. Because Lositan may show partially divergent results among independent simulations, we only considered candidate loci conservatively identified in three independent Lositan simulations assuming an infinite allele model of mutation, with a confidence interval of 0.99 and a FDR < 0.05, using the neutral mean F_ST_ and forcing the mean F_ST_ options.

Demographic histories can affect the statistical power of tests of selection (Lotterhos & Whitlock, 2015); thus, type I and type II errors are frequently associated with these approaches (Narum & Hess, 2011). We minimized the potential for false discovery by identifying consensus candidate loci for selection among distinct methods used for two datasets: (1) the entire genotypic dataset, including PA-arid individuals, using Lositan and Pcadapt consensus candidates and (2) a subset of genotypic data, which excluded the PA-arid samples due to absence of environmental data, using Lositan, Pcadapt and LEA consensus candidates (Supplemental Figure S2).

### 2.6 Functional annotation of candidate loci putatively under natural selection

Functional annotation of candidate loci was obtained by reciprocal nucleotide alignment between nextRAD sequences and the reference transcriptome characterized in the present study. Blast+ 2.2.31 (Camacho et al., 2009) was used with a threshold of at least 50 aligned nucleotides, a maximum of one mismatch and no gaps (Supplemental Figure S2).

### 2.7 Plant material for transcriptome assembly and differential expression analysis

On August 16^th^, 2013, three similarly sized seedlings of *A. germinans* with 5-9 stem nodes were collected from the PA-arid and PA-humid sites, separated by the Bragança-Ajuruteua road, and transplanted with the surrounding soil into 3.0-L pots (Figure 2c). Based on previous observations of other *Avicennia* species (Almahasheer, Duarte, & Irigoien, 2016; Duarte, Thampanya, Terrados, Geertz-Hansen, & Fortes, 1999), we estimate that all samples were between at least 3.5 months to at most 11 months old. For acclimation in a homogeneous environment, pots were placed in open-air and naturally shaded conditions and were watered daily after the sunset, with 300 mL of tap water. Plants were harvested at noon, after 72 hours of acclimation, washed with water and split into roots, stems and leaves with a sterile blade to be stored in RNAlater (Ambion Inc., Austin, TX, USA) and transported to the laboratory for RNA extraction.

### 2.8 RNA extraction, cDNA library preparation and RNA sequencing

RNA extraction was performed according to Oliveira, Viana, Reátegui, & Vincentz (2015). To assess purity, integrity and concentration, we used 1% agarose gel electrophoresis and a NanoVue spectrophotometer (GE Healthcare Life Sciences, Buckinghamshire, UK). Subsequently, cDNA-enriched libraries were constructed using TruSeq RNA Sample Preparation kits (Illumina Inc., California, USA). Libraries qualities were assessed using an Agilent 2100 Bioanalyzer (Agilent Technologies, California, USA) and concentrations were quantified using quantitative real-time PCR (qPCR), with a Sequencing Library qPCR Quantification kit (Illumina Inc.). Sequencing was performed with two 36-cycle TruSeq SBS paired-end kits (Illumina Inc.) on a Genome Analyzer IIx platform (Illumina Inc.).

### 2.9 Transcriptome assembly and functional annotation of transcripts

Raw data were filtered by quality, using Phred >20 for 70% of the read length, and adapters were trimmed using NGS QC Toolkit 2.3 (Patel & Jain, 2012). Filtered reads were *de novo* assembled into transcripts using the CLC Genomics Workbench (https://www.qiagenbioinformatics.com/). The distance between paired-end reads was set to 300-500 bp, the k-mer size was set to 45 bp, and the remaining default parameters were not changed.

Reads were mapped to the transcriptome using Bowtie1 (Langmead, Trapnell, Pop, & Salzberg, 2009), and contiguous sequences (contigs) without read-mapping support were removed from the assembly. For transcript annotation, we used blast+ 2.2.31 (Camacho et al., 2009), with an e-value <1e-10, using reference sequences from manually curated databases, as the National Center for Biotechnology Information (NCBI) RefSeq protein and RefSeq RNA (O’Leary et al., 2016) and representative proteins and cDNA from The Arabidopsis Information Resource (TAIR) (Berardini et al., 2015). We removed putative contaminant contigs from the assembly, which did not match plant sequences but showed high similarity to nonplant sequences from the NCBI’s RefSeq database. Sequences were assigned to the Kyoto Encyclopedia of Genes and Genomes (KEGG) orthology (KO) identifiers using the KEGG Automatic Annotation Server (KAAS) (Moriya, Itoh, Okuda, Yoshizawa, & Kanehisa, 2007). Protein family domains were searched using the Pfam database and the HMMER3 alignment tool (Finn et al., 2014).

We identified putative open reading frames (ORFs) using the program TransDecoder (http://transdecoder.sf.net) with default parameters. Redundant transcripts were detected using Cd-hit-est 4.6.1 (Li & Godzik, 2006) in the local alignment mode, with 95% identity and 70% coverage of the shortest sequence thresholds. To minimize redundancy, in the final assembly, we retained only sequences with the longest putative ORF and the longest putative non-coding transcripts from each Cd-hit-est clusters. Functional categories were assigned to putative coding sequences using the *Arabidopsis thaliana* association file from the Gene Ontology Consortium website (Blake et al., 2015) (Supplemental Figure S2).

### 2.10 Analysis of differentially expressed transcripts (DET)

Counts of reads mapped to assembled transcripts per sequenced sample were used as input files in DET analyses. Reads that mapped to multiple transcripts were excluded. The count matrix was normalized and used to detect transcripts with significant differential expression between PA-arid and PA-humid samples from the Bragança-Ajuruteua road (Figure 2) with the EdgeR Bioconductor package (Robinson, McCarthy, & Smyth, 2010) at a FDR <0.05. Gene Ontology (GO) term enrichment analyses were performed using the goseq R Bioconductor package (Young, Wakefield, Smyth, & Oshlack, 2010), which takes the length bias into account, with the default Wallenius approximation method and a p-value cutoff set to <0.05 (Supplemental Figure S2).

## 3. Results

### 3.1 Population genetics analyses of *A. germinans* along the equatorial coast of Brazil

Genetic diversity indices are shown in Table 2. The lowest levels of genetic variation were observed in the TMD site. The remaining sites, influenced by the northern branch of the SEC, presented considerably higher levels of diversity, including the PA-arid site, which was one of the most genetically diverse across the study region. All sampling sites deviated from Hardy-Weinberg equilibrium (HWE), with an excess of heterozygosity in all sites, besides ALC, which showed a small heterozygosity deficit (F_IS_=0.03). HWE deviation was highest in TMD (F_IS_=-0.43) but relatively low in all remaining sites, ranging from −0.15 (PNB) to +0.03 (ALC).

**Table 2.** Genetic diversity statistics, based on analyses of 2,297 genome-wide polymorphic loci detected in 57 individuals sampled in seven sampling sites along the equatorial Atlantic coastline of South America. π : Nucleotide diversity; H_E_: expected heterozygosity; H_O_: observed heterozygosity; pA: private alleles; %Poly: percentage of polymorphic loci; F_IS_ :inbreeding coefficient.

| Sampling site | $\pi$ (mean) | $\pi$ (s.e.) | $H_E$ | $H_O$ | pA | %Poly | $F_{IS}$ | Low Level (95% c.i.) | High Level (95% c.i.) |
| --- | --- | --- | --- | --- | --- | --- | --- | --- | --- |
| MRJ | 0.264 | 0.206 | 0.244 | 0.292 | 1 | 72.70 | -0.136 | -0.154 | -0.102 |
| PA-arid | 0.317 | 0.189 | 0.295 | 0.326 | 1 | 83.76 | -0.050 | -0.067 | -0.022 |
| PA-humid | 0.301 | 0.186 | 0.280 | 0.328 | 0 | 84.46 | -0.113 | -0.132 | -0.084 |
| ALC | 0.282 | 0.214 | 0.249 | 0.269 | 0 | 69.83 | 0.030 | 0.009 | 0.070 |
| PNB | 0.208 | 0.210 | 0.195 | 0.234 | 0 | 58.21 | -0.154 | -0.180 | -0.111 |
| PRC | 0.323 | 0.193 | 0.296 | 0.327 | 1 | 82.15 | -0.035 | -0.053 | -0.002 |
| TMD | 0.123 | 0.192 | 0.130 | 0.168 | 46 | 34.52 | -0.426 | -0.452 | -0.376 |

A substantial genetic structure was observed over the Equatorial Brazilian coast (Figure 3). The genetic diversity, based on 2,297 genome-wide SNPs genotyped in 57 individuals, was organized into four distinct genetic clusters (K=4) (Figure 3a, Supplemental Figure S3a-b). The greatest genetic divergences (F_ST_>0.46; Nei’s distance>0.255) were observed between individuals from TMD and those from all other sites (Figure 3b-c, Supplemental Table S1). The TMD site also presented the highest number of private alleles (pA=46). Most remarkable was the divergence between individuals from PA-arid and its adjacent site, PA-humid (F_ST_=0.25; Nei’s distance=0.181). This divergence was even greater than the observed divergence of PA-arid from PRC individuals (Supplemental Table S1), located approximately 900 km distant from each other (F_ST_=0.23; Nei’s distance=0.177), on the semi-arid coast of Brazil. The divergent gene pool of the PA-arid population was also evident when these samples were excluded from structure analyses. The most likely number of ancestral populations dropped from four to three (Figure 3c-d, Supplemental Figure S3c-d), and the overall structure remained unchanged (Figure 3d). At a finer scale, individuals from PA-humid and MRJ sites, on the Amazon Macrotidal Mangrove Coast (AMMC), seemed to be derived from the same ancestral population. This population, in turn, diverged from the population that may have given rise to individuals from ALC, PNB, and PRC sites, in Northeast Brazil (Figure 3a-b).

**Figure 3.**
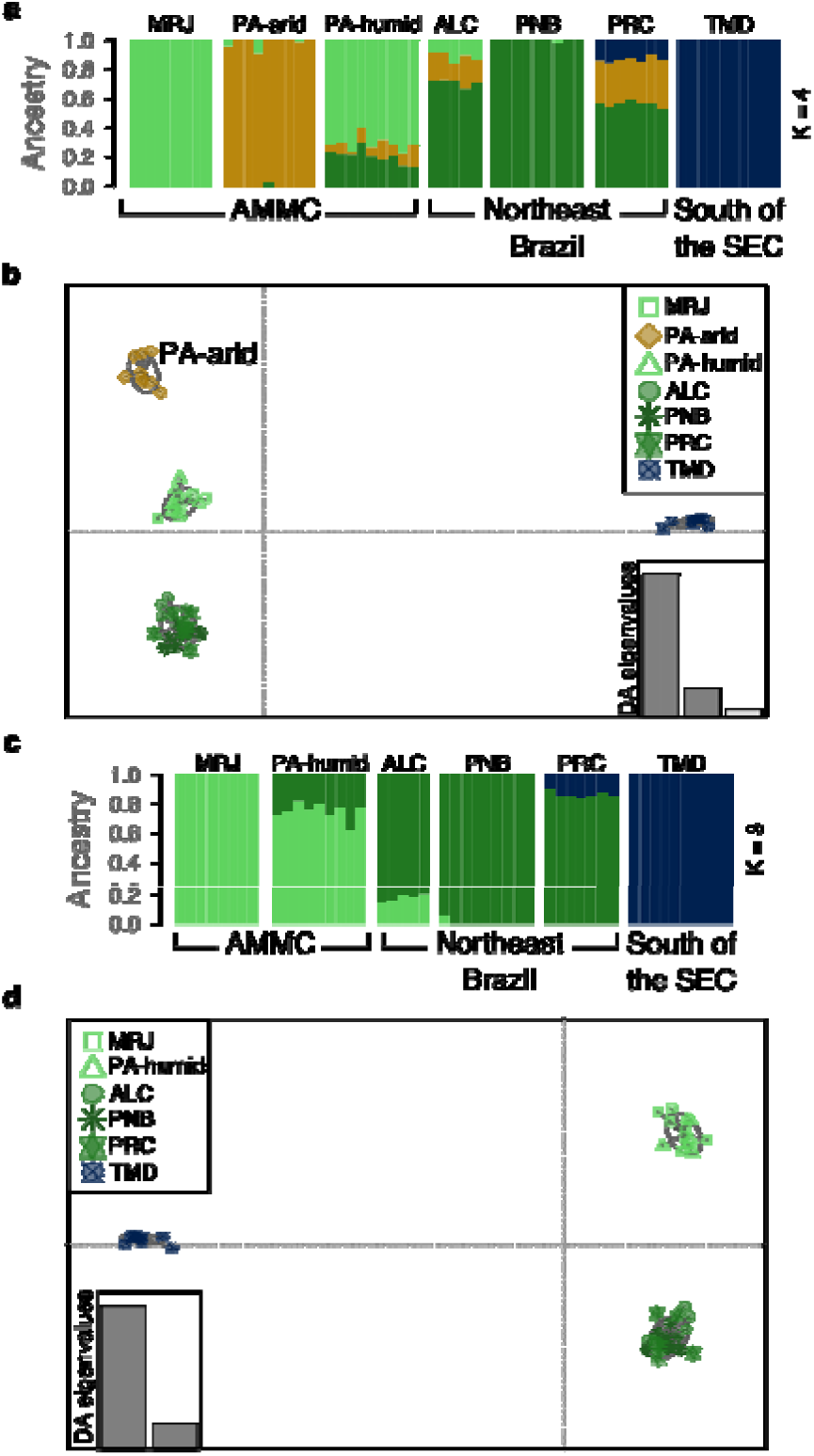
Genetic structure inferred from genome-wide single nucleotide polymorphisms (SNPs) detected in *Avicennia germinans*. (a) Attribution of ancestry implemented in the program Admixture 1.3.0 for all sampled individuals; stacked bars represent individuals, and each color represents one ancestral cluster (K=4). (b) Scatterplot of the first two principal components of the multivariate discriminant analysis of principal components (DAPC) of total genetic variance; all sampled individuals are represented as points; distinct symbols indicate sampling sites. (c) Attribution of ancestry (using Admixture 1.3.0) for all sampled individuals, excluding PA-arid samples; stacked bars represent individuals, and each color represents one ancestral cluster (K=3). (d) Scatterplot of the first two principal components of the DAPC of total genetic variance; sampled individuals, except for individuals sampled in the PA-arid site, are represented as points; distinct symbols indicate sampling sites.

### 3.2 *De novo* assembly and annotation of the *Avicennia germinans* reference transcriptome

We used RNA-Seq to sequence and *de novo* assemble a reference transcriptome from leaves, stems and roots of *A. germinans* seedlings, providing a functional context for this study. A total of 249,875,572 high-quality-filtered, paired-end, 72-bp reads, representing 78.25% of the raw data was used in the assembly. The reference transcriptome comprised 47,821 contigs, after removal of misassembled, redundant and contaminant sequences. Putative ORFs were identified in 29,854 contigs, subsequently annotated as putative protein-coding transcripts. The remaining 17,967 contigs were classified as putative non-coding transcripts. A detailed characterization of contigs can be found in Supplemental Table S2.

Raw reads were mapped back to the reference transcriptome, with over 82% uniquely mapped to a single transcript and only 1.31% mapped to more than one transcript (Supplemental Table S3). We found 91.74% of the plant universal orthologs from the BUSCO database (Simão, Waterhouse, Ioannidis, Kriventseva, & Zdobnov, 2015) represented in the reference transcriptome (Supplemental Table S3). We also found from 20,529 to 31,348 putative orthologous sequences between the *A. germinans* transcriptome and four other publicly available transcriptomes derived from the genus *Avicennia* L. (Acanthaceae) (Supplemental Table S4).

As expected, most putative protein-coding transcripts of the reference transcriptome could be annotated using relevant databases (92.6%), whereas only a few putative non-coding transcripts could be annotated (33.6%) (Supplemental Figure S4). We found 2,207 putative coding transcripts (7.4%) and 11,925 putative non-coding transcripts (66.4%) unique to *A. germinans*, which may represent lineage specific sequences.

### 3.3 Detection of candidate loci responding to environmental selection

Genome-wide signatures of selection were detected from genotypic data retrieved from samples from all seven sites of collection. Fifty-six putative outlier loci were consistently identified by two F_ST_ outlier methods, solely based on deviations from neutral expectations of the distribution of genetic diversity. Eleven of these loci aligned to sequences in the reference transcriptome, of which eight were highly similar to proteins associated with the response or tolerance to drought in *A. thaliana* or *Sesamum indicum* (Figure 4a, Supplemental Figure S5).

**Figure 4.**
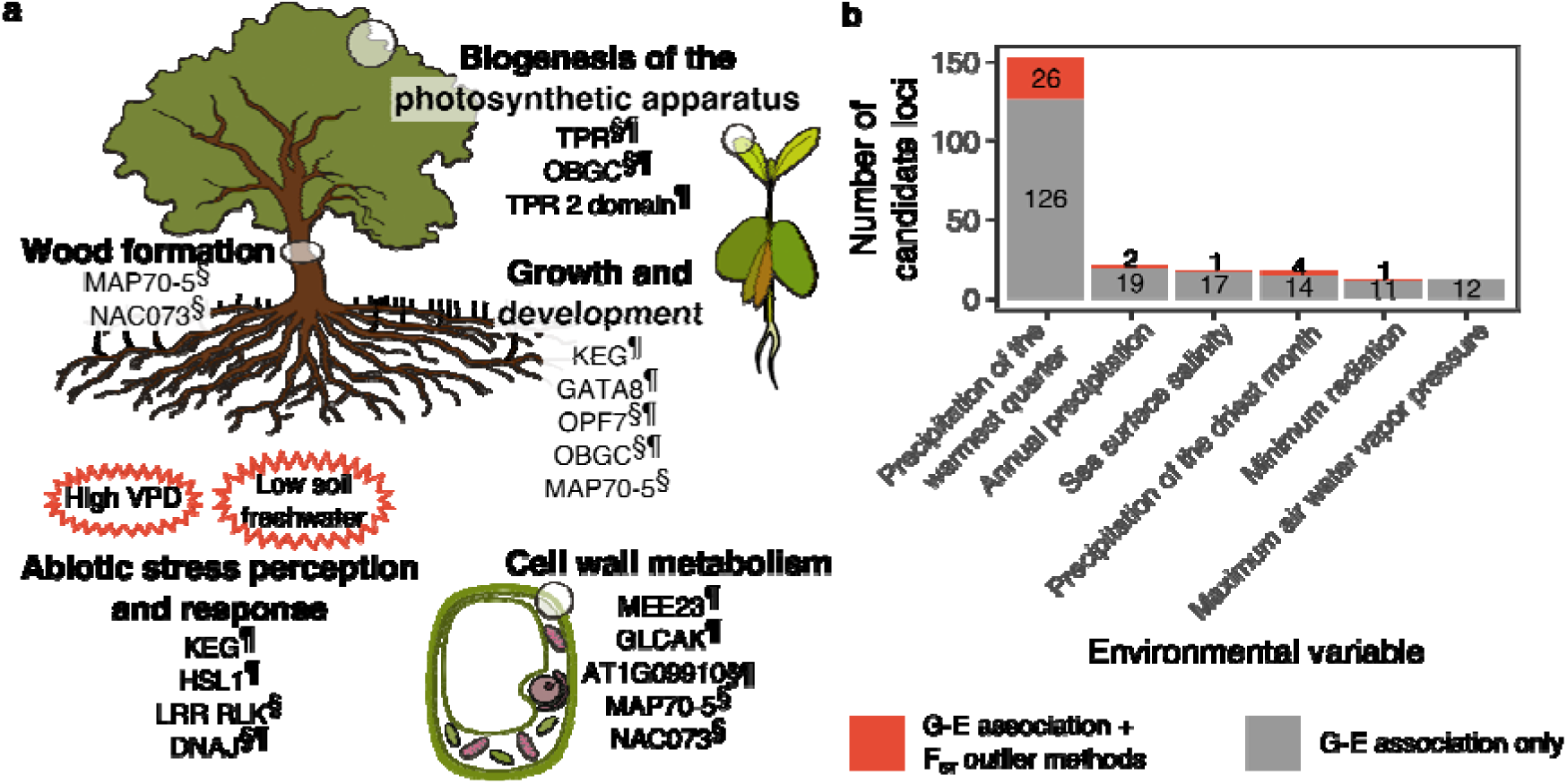
Candidate loci for selection detected from *Avicennia germinans* sampled along tropical mangrove forests of the north-northeastern Brazilian coast. (a) Schematic representation of key biological processes associated with candidate loci. *Arabidopsis thaliana* gene symbols listed below biological processes represent the functional annotation of candidate loci present within putative protein-coding regions of the genome. *§ Labeled gene symbols*: candidates obtained using F_ST_ outlier tests performed with all sampled individuals, including PA-arid samples. *¶ Labeled gene symbols*: candidates obtained by combining genetic-environmental association tests and F_ST_ outlier approaches, using a subset of samples, without PA-arid ones, for which environmental characterization could not be retrieved from public databases. (b) Number of candidate SNPs associated with non-collinear environmental variables.

The exclusion of PA-arid samples was necessary for G-E association tests, due to limitations in the resolution of Marspec and WorldClim environmental layers. From the remaining subset of 48 individuals from six sampling sites, further 153 candidate loci for selection were identified by G-E correlation and two F_ST_ outlier approaches (Figure 4b). Out of these candidate loci, 24 aligned to the reference transcriptome, of which 20 were putative protein-coding showing high similarity to gene models from *A. thaliana* or *S. indicum* (Supplemental Figure S5).

Among all candidate loci for selection detected along the sampling region, we found 14 loci associated with plant growth and development, wood formation, cell wall metabolism, biogenesis of the photosynthetic apparatus, abiotic stress perception and response and protein protection from stress-induced aggregation, among other processes (Figure 4a, Table 3).

**Table 3.** Annotation of candidate SNP loci putatively under selection effect identified by two (Lositan and Pcadapt) or three (Lositan, Pcadapt and LEA) distinct methods, from two datasets. (1) All sampled individuals, including PA-arid samples, using F_ST_ outlier approaches and (2) a subset of samples, excluding individuals from the PA-arid site, combining genetic-environmental association tests and F_ST_ outlier tests.

| Dataset | Transcript ID | Similar to† | Environmental association‡ | Putative function |
| --- | --- | --- | --- | --- |
| 1 | Ag_23357 | AT4G28500, NAC073, NAC domain containing protein 73 | NA | Regulation of the biosynthesis of cellulose and hemicellulose in wood fibers and the expression of lignin-polymerizing and signaling genes (Hussey et al., 2011; Zhong, Lee, Zhou, McCarthy, & Ye, 2008) |
| 1 | Ag_47094 | AT4G17220, MAP70-5, microtubule-associated proteins 70-5 | NA | Fundamental for the development of secondary cell wall band patterning in xylem tracheids and for wood formation, playing a role in the anisotropic expansion of cells (Pesquet, Korolev, Calder, & Lloyd, 2010) |
| 1 | Ag_26495 | AT5G67200, Leucine-rich repeat protein kinase (LRR-RLK) family protein | NA | May play a role in hormones and abiotic stress sensing and signaling (ten Hove et al., 2011; Van der Does et al., 2017) and in regulating adaptation to salt stress (Lorenzo et al., 2009) |
| 1 | Ag_43440 | Not annotated | NA | Not annotated |
| 1 and 2 | Ag_4919 | AT1G09910, Rhamnogalacturonate lyase family protein | PWQ (8.31e-05) | Associated with the degradation of the cell wall polysaccharides, and pectin (McDonough, Kadirvelraj, Harris, Poulsen, & Larsen, 2004) |
| 1 and 2 | Ag_20543 | AT5G18950, Tetratricopeptide repeat (TPR)-like superfamily protein | PWQ (0.00018) | Associated with photosynthetic machinery biogenesis, stabilization and repair (Bohne, Schwenkert, Grimm, & Nickelsen, 2016) |
| 1 and 2 | Ag_5627 | PREDICTED: GTP-binding protein OBGC, chloroplastic [Sesamum indicum] | PWQ (8.31e-05) | Associated with thylakoid membrane biogenesis and chloroplast protein synthesis and essential for early embryogenesis in response to light stimulus (Bang et al., 2009) |
| 1 and 2 | Ag_29619 | AT2G18500, OFP7, ovate family protein 7 | PWQ (0.00019) | May be involved in various aspects of plant growth and participate in suppressing cell elongation (Wang, Chang, & Ellis, 2016; Wang, Chang, Guo, & Chen, 2007) |
| 1 and 2 | Ag_38780 | AT2G24395, chaperone protein dnaJ-related | PWQ (8.31e-05) | Regulation of the activity of Hsp70 chaperones (P. Walsh, Bursac, Law, Cyr, & Lithgow, 2004) and protein protection against stress-induced denaturation (Frydman, 2001) |
| 2 | Ag_12760 | AT3G54810, GATA8, Plant-specific GATA-type zinc finger transcription factor family protein | PWQ (0.00363); PDM (0.01916) | Associated with the control of plant growth and development (Manfield, Devlin, Jen, Westhead, & Gilmartin, 2006) |
| 2 | Ag_135 | AT1G28440, HSL1, HAESA-like 1 | PWQ (9.09e-05) | The Ag_135 transcript has a kinase domain and is associated with growth and abiotic stress signaling (ten Hove et al., 2011) |
| 2 | Ag_15249 | AT2G34790, MEE23, FAD-binding Berberine family protein | PWQ (8.31e-05) | Involved in the lignification of the cell wall; mediates the oxidation of monolignols (Daniel et al., 2015) and is required for endosperm development (Pagnussat et al., 2005) |
| 2 | Ag_10980 | AT5G13530, KEG, keep on going | PWQ (0.00041); PDM (0.02034) | Associated with the control of plant growth and development and critical for seedling regulation of abscisic acid perception and signaling (Stone, Williams, Farmer, Vierstra, & Callis, 2006) |
| 2 | Ag_26982 | AT3G01640, GLCAK, glucuronokinase G | PWQ (0.00011) | Involved in the biosynthesis of UDP-glucuronic acid, providing nucleotide sugars for the polymerization of cell-wall compounds (Pieslinger, Hoepflinger, & Tenhaken, 2010) |
| 2 | Ag_11269 | AT2G25730, unknown protein | PWQ (0.00016) | The Ag_11269 transcript has a TPR 2 domain and, thus, may be involved in thylakoid membrane biogenesis and with the photosynthetic machinery stabilization and repair (Bohne, Schwenkert, Grimm, & Nickelsen, 2016) |
| 2 | Ag_43282 | AT1G04300, MUSE13, TRAF-like superfamily protein | PWQ (8.31e-05) | MUSE13 regulates the turnover of nucleotide-binding domain and leucine-rich repeat-containing immune receptors. Loss of MUSE13 and MUSE14 leads to enhanced pathogen resistance, NLR accumulation, and autoimmunity. May regulate ATG6 ubiquitination and formation of autophagosomes (Huang et al., 2016) |
| 2 | Ag_14879 | AT2G42520, P-loop containing nucleoside triphosphate hydrolases superfamily protein | PWQ (0.00109) | AT2G42520 has important roles in RNA metabolism such as splicing, RNA transport, ribosome biogenesis, translation and decay and unwinds double-stranded RNA molecules through the hydrolysis of NTP |
| 2 | Ag_20289 | AT1G48650, DEA(D/H)-box RNA helicase family protein | PDM (0.01222); PANNUAL (5.80E-05); SSS (0.01461); WVPMAX (0.03580); RADMIN (0.03813) | AT1G48650 has important roles in RNA metabolism such as splicing, transport, ribosome biogenesis, translation and decay; it unwinds double-stranded RNA molecules through the hydrolysis of NTP |
| 2 | Ag_24006 | AT1G11050, Protein kinase superfamily protein | PWQ (0.00341); PDM (0.00671); PANNUAL (0.01962) | Ag_24006 has a protein kinase domain |
| 2 | Ag_22233 | AT5G01020 Protein kinase superfamily protein | PWQ (0.00022) | Ag_22233 has a protein kinase domain |
| 2 | Ag_43463 | AT5G43960, Nuclear transport factor 2 (NTF2) family protein with RNA binding (RRM-RBD-RNP motifs) domain | PWQ (0.00011) | mRNA-binding protein, controls the fate and expression of a transcript |
| 2 | Ag_36772 | AT2G35680, Phosphotyrosine protein phosphatases superfamily protein | PWQ (0.00026) | AT2G35680 may possess phosphatase activity |
| 2 | Ag_13254 | AT4G30100, P-loop containing nucleoside triphosphate hydrolases superfamily protein | PWQ (8.31e-05) | AT4G30100 has a hydrolase activity |
| 2 | Ag_29952 | AT5G11970,<br>Protein of unknown<br>function (DUF3511) | PWQ<br>(8.62e-05) | Unknown function |
| 2 | Ag_1345 | AT5G55960,<br>unknown protein | PWQ<br>(0.00018) | Unknown function |
| 2 | Ag_43837 | Not annotated | PWQ<br>(8.62e-05) | Not annotated |

†Blastx hit to *Arabidopsis thaliana* or *Sesamum indicum* [when indicated] gene models. ‡(*FDR*): False discovery rate values (Benjamini-Hochberg procedure) for genetic-environment association tests; *NA*: genetic-environment association tests were not available for dataset (1) due to unavailability of environmental data for the PA-arid sites in public databases; *PWQ*: Precipitation of the warmest quarter; *PDM*: Precipitation of the driest month; *PANNUAL*: Annual precipitation; *SSS*: Mean sea surface salinity; *WVPMAX*: Maximum water vapor pressure; *RADMIN*: Minimum radiation.

### 3.4 Differential transcript expression analysis

Transcriptome sequencing of seedlings grown under contrasting field conditions, revealed significant expression differences in 2,454 transcripts, despite previous acclimatization under homogeneous shaded, well-watered conditions (Figure 2c). Most DETs were detected in roots (2,337) and stems (1,383), followed by leaves (361) (Figure 5, Supplemental Figure S6). We refer to DETs that showed higher expression levels in samples from the PA-arid site than in samples from the PA-humid site as “DET-Arid” and to DET showing a significantly higher expression in samples from the PA-humid than in samples from the PA-arid site as “DET-Humid”.

**Figure 5.**
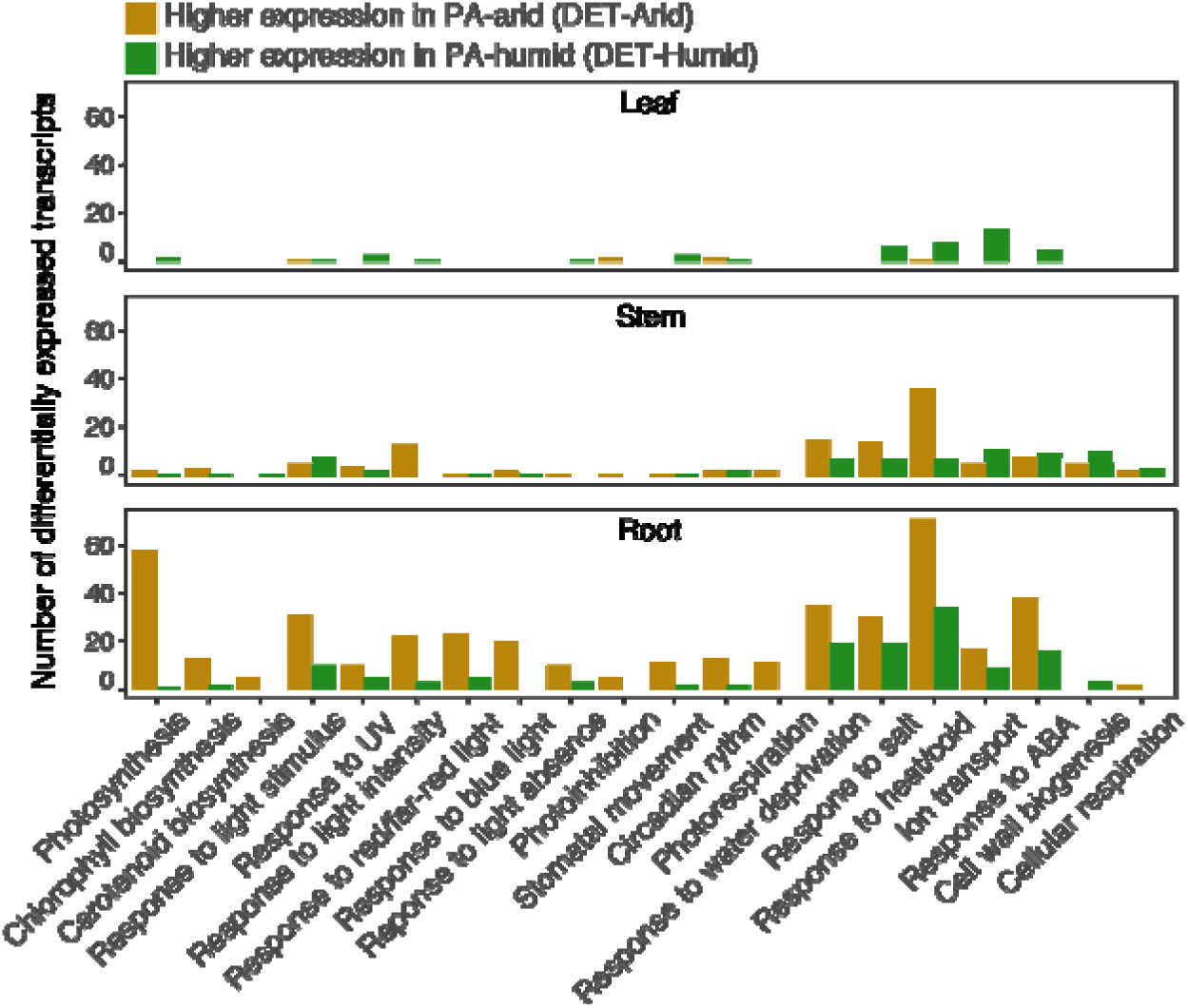
Functional categories of differentially expressed transcripts (DETs) identified between samples grown in the PA-arid and PA-humid sampling sites. Categories were selected based on biological processes previously identified to be involved in the response, acclimation or resistance to water stress in model plants.

The functional annotation and subsequent assignment of most putative protein-coding transcripts to GO terms (76.24%) (Supplemental Figure S4) enabled the assessment of DETs that highlighted key aspects of a differential response of *A. germinans* to contrasting source environments, differing markedly in hydrological regime, soil pore water salinity, solar radiation and surface temperature (Lara & Cohen, 2006; Pranchai et al., 2017; Vogt et al., 2014). We focused this analysis on enriched biological processes previously identified to be involved in the tolerance, resistance or response to osmotic and drought stress in various crops, model and non-model species, including mangroves (Figure 5). A detailed characterization of the differentially transcripts belonging to such processes is shown in Supplemental Table S5. In the following subsections we summarize these results.

#### 3.4.1. Photosynthesis

Mangroves have aerial, photosynthesizing roots, which contribute to carbon gain and enable root respiration in anaerobic soils, using both atmospheric and photosynthetically regenerated oxygen (Kitaya et al., 2002). Therefore, although various photosynthesis genes frequently express only in basal levels in roots of the model plant, *Arabidopsis thaliana* (Klepikova, Kasianov, Gerasimov, Logacheva, & Penin, 2016), they may express in higher levels in mangroves, even during the seedling stage, when aerial roots are generally absent, as observed in *A. germinans*, and in the congeneric *A. officinalis* (Krishnamurthy et al., 2017). Differential transcripts expression analyses revealed that photosynthesis-associated transcripts were enriched, mainly in roots and stems from the DET-Arid set. These included putative enzymes and genes required for chloroplast biogenesis or development, for chlorophyll biosynthesis, for the assembly of the photosynthetic apparatus and light-harvesting complexes and involved in acclimation to fluctuating light. The DET-Arid set of roots samples also presented transcripts associated with the C3 carbon fixation pathway, including, glyceraldehyde-3-phosphate dehydrogenase subunits, calvin cycle proteins, and the RIBULOSE BISPHOSPHATE CARBOXYLASE SMALL CHAIN 1A (RBCS1A), which showed a 7.8 Fold-Change relative to roots of PA-humid samples.

#### 3.4.2. Response to light

The DET-Arid sets of roots and stems were enriched in transcripts associated with the response to light but not directly associated with photosynthesis. These included transcripts similar to genes and transcription factors involved with light acclimation and essential for maintaining electron flow and photosynthetic efficiency under changing light conditions (Privat, Hakimi, Buhot, Favory, & Lerbs-Mache, 2003). The DET-Arid sets of roots and stems also included putative light-signaling genes and photoreceptors, as phototropins, cryptochromes and phytochromes. These photoreceptors are sensitive to light intensity and control complex light and stress responses, including photoinduced movements as well as growth and development under limiting light (Correll et al., 2003; Ohgishi, Saji, Okada, & Sakai, 2004; Pedmale et al., 2016). Complementarily, we identified transcripts similar to proteins that interact with these photoreceptors in the mediation of shade avoidance and phototropism under low light, and to low light-induced transcription factors, which regulate developmental processes and growth in response to shade avoidance. Moreover, in the DET-Arid sets of roots and stems, we found transcripts associated with chloroplast accumulation upon low blue light and with the response to sugar starvation induced by dark.

#### 3.4.3. Response to water deprivation and response to salt

The DET-Arid sets of roots and stems were also enriched in transcripts associated with the response to osmotic stress and water deprivation, including putative genes that play relevant roles in drought and salt stress resistance. For instance, we found transcripts associated with abscisic acid (ABA)-dependent stomatal closure, a well-known mechanism for maintaining water status under drought, which severely compromises growth (Murata, Mori, & Munemasa, 2015). Additionally, DET-Arid sets of roots and stems included transcripts associated with drought-induced ion transporters, which protect chloroplasts from deleterious Na^+^ concentrations (Müller et al., 2014) and involved in Ca^2+^ homeostasis under abiotic stress. Complementarily, we found in the DET-Arid sets of roots and stems transcripts associated with biosynthesis and accumulation of oligosaccharides, which may act as osmoprotectants, increasing tolerance to osmotic stress (ElSayed, Rafudeen, & Golldack, 2014; Nishizawa, Yabuta, & Shigeoka, 2008; Taji et al., 2002). Additionally, transcripts associated with drought-induced epicuticular wax biosynthesis, transport and deposition in cell walls were among DET-Arid sets of roots and stems. We also detected in these sets of DETs several transcripts associated with reactive oxygen species (ROS) dissipation for the control of cell damage caused by drought, high light or salt stress, including copper/zinc superoxide dismutases, a chloroplastic drought-induced stress protein (CDSP32) and MAP KINASE 6 (MAPK6).

#### 3.4.4. Response to heat

Transcripts similar to genes that confer tolerance to heat were enriched among the DET-Arid sets of roots and stems. We detected several putative co-chaperones and chaperones, which protect proteins from structural degradation (Al-Whaibi, 2011), and transcripts associated with putative components of thermomemory, required for long-term maintenance of acquired thermotolerance (Charng et al., 2006).

#### 3.4.5. Photorespiration

As a response to drought, plants close their stomata, reducing the CO_2_:O_2_ ratio in mesophyll cells, increasing levels of photorespiration. Rates of photorespiration may also be increased by high temperatures, which reduce both the solubility of CO_2_ in the mesophyll cells and the ability of the ribulose-1,5-bisphosphate carboxylase/oxygenase (RuBisCO) to discriminate between O_2_ and CO_2_ (Kozaki & Takeba, 1996; Wingler, Lea, Quick, & Leegood, 2000). On the one hand, photorespiration decreases photosynthesis efficiency, on the other, it plays an essential role in protecting the photosynthetic machinery from damage caused by excessive photochemical energy (Kozaki & Takeba, 1996; Wingler et al., 2000). In the DET-Arid set of roots, we detected enrichment of putative genes associated with photorespiration, similar to enzymes participating in transamination, required for maintaining photosynthesis under photorespiratory conditions and for carbon flow during photorespiration, and which catalyze the concluding reaction of photorespiration.

## 4. Discussion

Our study suggests that spatial variation in freshwater availability play an important role in driving adaptation of the typically tropical and widespread tree, *Avicennia germinans*. In the following subsections, we describe how variation in oceanographic and bioclimatic variables along the equatorial Brazilian coast seem to influence the organization of genome-wide genetic diversity of the species. Moreover, we show that drastic and persistent restriction in soil freshwater availability likely caused the rapid evolution of phenotype, gene pool and gene expression profile of a recently founded population of *A. germinans*, without depletion of genetic diversity, and despite clear possibility of gene flow with an adjacent, unchanged population.

### 4.1 Gradual environmental variation in freshwater availability may partly explain the organization of non-neutral genetic variation in *A. germinans*

The genetic structure inferred by genome-wide SNPs (Figure 3) suggested the importance of both neutral and non-neutral environmental drivers of variation. The greatest divergence was observed between samples influenced by the northern and southern branches of the South Equatorial sea current (SEC) (Figure 1), acting on the Atlantic coast of South America, corroborating previous results found for other coastal trees, using putatively neutral molecular markers (Francisco, Mori, Alves, Tambarussi, & Souza, 2018; Mori, Zucchi, & Souza, 2015; Takayama, Tateishi, Murata, & Kajita, 2008). The flow direction of coastal currents provides a neutral explanation to the structure of genome-wide diversity of sea dispersed species, due to restrictions in the dispersal of buoyant propagules, likely facilitating the accumulation of random genetic divergences (Francisco et al., 2018; Mori, Zucchi, & Souza, 2015; Wee et al., 2014; Yan, Duke, & Sun, 2016). Our results also suggest that the lower precipitation of the warmest quarter in northern sites (AMMC and Northeast Brazil regions) (Figure 1) may play an additional, non-neutral role in shaping this north-south divergence. We hypothesize that the more even distribution of rainfall throughout the year in the TMD site likely alleviates water-stress in *A. germinans*, whereas limited rainfall in the warmest quarter of northern regions increases soil aridity and salinity, reducing opportunities for rehydration and potentially favoring increased drought-tolerance. It is plausible that this environmental filter contributes to the genetic divergence observed between TMD and remaining populations (Figure 3). This hypothesis is corroborated by the detection of 26 loci candidate for selection correlated to the spatial variation in the precipitation of the warmest quarter over the study area (Figure 4b). Although most of these loci were poorly characterized, hampering inferences about their functional relevance in the environmental context, we were able to associate 11 candidates with biological processes influenced by drought, as photosynthesis, cell wall metabolism, cell elongation, plant growth, protein protection from stress-induced degradation and regulation of abscisic acid signaling (Figure 4, Table 3). The adaptive importance of freshwater limitation in tropical trees was also suggested in the mangrove, *A. schaueriana* (M. V. Cruz et al., 2018), and in tropical forests and savannas (Ciemer et al., 2019), for which it was similarly suggested that an environmental filtering mechanism driven by rainfall variability likely favored the survival of more drought resistant lineages.

Although *Avicennia* propagules can remain viable for long periods and present transoceanic dispersal (Mori, Zucchi, Sampaio, et al., 2015), at a finer scale, we observed a genetic divergence between sites on the AMMC region (MRJ and PA-humid) and on Northeast Brazilian mangroves (ALC, PNB and PRC) (Figure 3). The AMMC region shows higher annual precipitation and is more strongly influenced by riverine freshwater inputs than the remaining sites, due to its closer proximity to the Amazon River Delta (Figure 1). Conversely, the Northeast Brazilian coastline is characterized by reduced rainfall and the lack of riverine freshwater inputs (Figure 1), which limit plants access to soil freshwater, due to increased salinity, potentially contributing to the local specialization of individuals. In fact, we were able to detect two loci associated with sea surface salinity data and total annual precipitation, based on both F_ST_ outlier methods and G-E association tests (Figure 4b). These loci showed similarity to a poorly characterized protein kinase and to an RNA hydrolase (Table 3). Given the unclear functional relevance of these putative adaptive loci in the environmental context, we recommend future efforts to analyze the role of demographic history in *A. germinans*, to find additional explanations for the genetic structure observed along the equatorial Brazilian coastline (Figure 3). Additionally, candidate loci detected in our study may play a role in drought adaptations, but their molecular functions need to be further characterized in plants adapted to physiological drought. The functional role of these proteins in drought tolerance may not get highlighted in genetic screenings performed on drought-sensitive model plants, distantly related to *A. germinans*.

### 4.2 Rapid evolution of *A. germinans* in response to abrupt limitation in access to soil freshwater

The PA-arid population of *A. germinans* was originated after 1974, when the construction of the Bragança-Ajuruteua road altered the hydrology of part of the mangrove forest in the AMMC region (Figure 2a-b) (Cohen & Lara, 2003). This sudden environmental change caused a large dieback of the mangrove vegetation, followed by a gradual recolonization, mainly by *A. germinans*. We estimate that the time since this event occurred, could represent at most 40 reproductive cycles of *A. germinans*, based on previous observations of closely related *Avicennia* sp. (Almahasheer et al., 2016; M. V. Cruz et al., 2018), although a conservative estimate could be four generations (Polidoro et al., 2010). The modification of the architecture of recolonizing individuals of PA-arid, from previous tall/arboreal, to now dwarf/shrub (Figure 2b) (Pranchai et al., 2017), suggests that limitation in plant access to soil water favored smaller tree sizes, one of the most integrative characteristic of drought resistance (Bennett et al., 2015; Corlett, 2016; Rowland et al., 2015). On the one hand, we cannot determine how much of the observed difference in tree size is inherited or determined by phenotypic plasticity. On the other hand, our results revealed a substantial change in genome-wide allele frequencies of trees, which likely represent samples of the first generations of the recolonizing population (Figure 3a, Table 2). Interestingly, changes in allele frequencies at founding occurred without reductions in levels of diversity, as estimated by various population genetics parameters (Table 2). Moreover, consistent with an adaptive response to selection, likely driven by restrictions in plants access to soil freshwater, F_ST_ outlier tests detected 56 candidate loci, of which eight were present within transcripts associated with functions as suppression of cell elongation, wood and xylem tracheids formation, photosynthetic machinery biogenesis and repair, regulation of adaptation to stress and of protection from stress-induced protein degradation (Figure 4a, Table 3). The simultaneous presence of high levels of diversity in the founding population (Table 2) and strong selection on individual traits is common in natural populations, and is a signature of substantial multivariate genetic constraints (B. Walsh & Blows, 2009), as expected in this specific case, given the magnitude of environmental change.

A surprising aspect of the population genetic analyses, however, was that genome-wide changes in allele frequencies occurred despite expectations of high levels of gene flow, given the proximity (<10.0 m) of contrasting populations, and characteristics of the species, which is preferably allogamous, with entomophyllic flowers (Mori, Zucchi, & Souza, 2015; Nadia et al., 2013; Nettel-Hernanz et al., 2013). In the absence of a geographic barrier to gene flow, recombination is expected to hinder adaptation, unless there is very strong selection or reproductive isolation (Kawecki & Ebert, 2004; Mayr, 1963). As we expect selection to be very strong in this case, due to extreme environmental change, it could be enough to counteract the homogenizing effect of gene flow or even to cause shifts in the timing of flowering (Cho, Yoon, & An, 2017), resulting in reproductive isolation.

Besides genetic changes, we also identified transcripts expression differences between seedlings from PA-arid and PA-humid sites after acclimatization in pots under shaded, well-watered conditions (Figure 2c, Figure 5), indicating an additional molecular basis for phenotypic divergences. Because differential gene expression influence trait variation (Wolf, Lindell, & Backstrom, 2010), it can be substantial between distinct locally adapted populations (Akman, Carlson, Holsinger, & Latimer, 2016; Gould, Chen, & Lowry, 2018). Differential expression was observed mainly in roots and stems of *A. germinans* seedlings, which are the first organs exposed to increased salinity and water deficits in mangroves, thus where osmotic stress sensing and signaling to the whole plant are triggered (Chaves, Maroco, & Pereira, 2003; Janiak, Kwas niewski, & Szarejko, 2016). Such transcriptome changes were associated with various biological processes previously identified to be involved in central aspects of osmotic-, heat- and UV-stress in model and non-model plants (Ding et al., 2013; Fan et al., 2018; Zhang et al., 2015). Being sessile under extreme environmental conditions, PA-arid plants should need to rapidly adjust photosynthesis gene expression to intermittent freshwater availability (Chaves, Flexas, & Pinheiro, 2009; Urban, Aarrouf, & Bidel, 2017). The increased freshwater supply and the restriction in solar irradiance under experimental conditions, compared to source-site (Figure 2c), likely required PA-arid individuals to broadly adjust their photosynthetic machinery. Conversely, although environmental conditions were relaxed in the RNA-Seq experiment, we observed several transcripts associated with high temperature, drought and salinity response among DET-Arid, including epicuticular wax and cutin synthesis, export and deposition, accumulation of osmoprotectants and antioxidants and ion homeostasis. We suggest that these findings represent key regulatory mechanisms, by which PA-arid plants tolerate a predominantly warm, highly saline and dry environment, with extreme UV levels. These strategies may have enhanced the relative capacity of PA-arid seedlings to grow under a high leaf water-deficit status and infrequent freshwater inputs.

We suggest that future efforts to understand the molecular mechanisms of adaptation of the PA-arid population should focus on characterizing epigenetic differences between PA-arid and PA-humid. Epigenetic changes can emerge faster than adaptive genetic changes, also playing an important role in early adaptive process (Kenkel & Matz, 2016; Pavey, Nosil, & Rogers, 2010). Although the mechanisms are not entirely clear, our results suggest, through independent molecular approaches, that a rapid evolution of this population occurred, with gene expression and genetic changes. The desertification of the PA-arid site represents a drastic and persistent environmental change, analogous to a freshwater exclusion experiment for mangroves of the tropical Brazilian coastline, in which this limitation likely caused the rapid evolution of recolonizing *A. germinans* individuals. Although it is difficult to demonstrate rapid evolution in nature, there is growing empirical evidence that when environmental selection is very intense, evolutionary processes may occur on a very fast time scale (Amorim et al., 2017; Donihue et al., 2018; Schoener, 2011), as observed in this case.

### 4.3 Implications for conservation

Wetlands of the Northeastern coast of Brazil and the ecosystem services they provide are predicted to be particularly impacted by future reductions in precipitation and freshwater availability (Osland et al., 2018). Despite these threats, the extant gene pool of northern populations of *A. germinans* seems to harbor sufficient diversity to enable species persistence through natural selection of drought-resistant plants, as observed in the case of PA-arid. Recolonizing individuals in this location share alleles with individuals from distant areas, as the PRC and ALC sites (Figure 3a), in Northeast Brazil, suggesting that these forests might have contributed as source of adaptive variation through sea-dispersed propagules in the recolonization of the PA-arid site, in addition to the selection on migrants from adjacent areas.

In the context of an increasingly drying climate, reforestation plans for populations of *A. germinans* located south of the SEC should consider using mixed stocks of seedlings (Moritz, 1999), to introduce genetic variation associated with increased drought-tolerance present in Northeastern mangroves, while also maintaining local alleles, possibly associated with site-specific environmental characteristics.

### 4.4 Concluding remarks

We provide novel insights into how limited access to soil freshwater can change allele frequencies, gene expression and phenotypes of a dominant tropical tree. These findings are consistent with the predictions of a hypothesis of natural selection, but we acknowledge that other evolutionary processes could also play a role in driving the observed patterns, opening new opportunities for future investigation. Research on the genomic basis of tree adaptation are often limited by difficulties in implementing empirical tests, given their long generation time and the scarcity of basic biological information. In tropical forests, mostly found in developing countries, the lack of resources imposes additional challenges for adaptation studies. Advances in the understanding of the genomic basis of drought-tolerance in tropical trees can support effective protection plans and mitigating climate change. As shown in this study, such knowledge can improve predictions of the persistence of the ecosystems they form and services they provide and generate key insight for conservation and management efforts (Holliday et al., 2017; Moran, Hartig, & Bell, 2016).

## Acknowledgments

We acknowledge Ilmarina Menezes for assistance with fieldwork and HPC services at Louisiana State University. M.V.C. received fellowships from the São Paulo Research Foundation - FAPESP (PhD 2013/26793-7) and from the Coordination for the Improvement of Higher Education Personnel - CAPES Computational Biology Program (PhD SWE 8084/2015-07, PhD 88887.177158/2018-00). G.M.M. received fellowships from FAPESP (PD 2013/08086-1, PD BEPE 2014/22821-9) and a grant from the Brazilian National Council for Scientific and Technological Development - CNPq (PD 448286/2014-9). M.D. and D-H.O. received grants from the United States National Science Foundation award (MCB 1616827) and the Next-Generation BioGreen21 Program of Republic of Korea (PJ01317301). R.S.O. received a CNPq productivity scholarship and a grant from Microsoft research-FAPESP (2011/52072-0). A.P.S. received a grant from CAPES - Computational Biology Program (88882.160095/2013-01), a research fellowship from CNPq (309661/2014-5) and a CNPq productivity scholarship.

## Conflicts of interest

The authors have no conflict of interest to disclose.

## Data accessibility

- Gene expression data and transcriptome sequences that support conclusions have been deposited in GenBank with the accession code GSE123659;
- SNP genotypes are available at Dryad doi:10.5061/dryad.h11t255 (M. V. Cruz et al., 2019).

## Author contributions

A.P.S., G.M.M. and M.V.C. designed the study. M.V.C. and G.M.M. conducted fieldwork, prepared samples for sequencing and wrote the manuscript. M.V.C., G.M.M., R.S.O., M.D. and D.H.O. analyzed the RNA-Seq and nextRAD results. A.P.S, M.I.Z., G.M.M. contributed material/reagents/analytical tools. All authors discussed the results and contributed to the manuscript.

## Notes

#### Summary of Updates

Major revisions of the entire manuscript text and figures were performed, following peer-reviewers suggestions and critiques. New genetic diversity analyses were made to better support our findings.

