## Supplemental for "Molecular responses to freshwater limitation in the mangrove tree *Avicennia germinans* (Acanthaceae)"

**Number of Supplemental Figures:** 6 (colored)

**Number of Supplemental Tables:** 5

**1.	Figures**

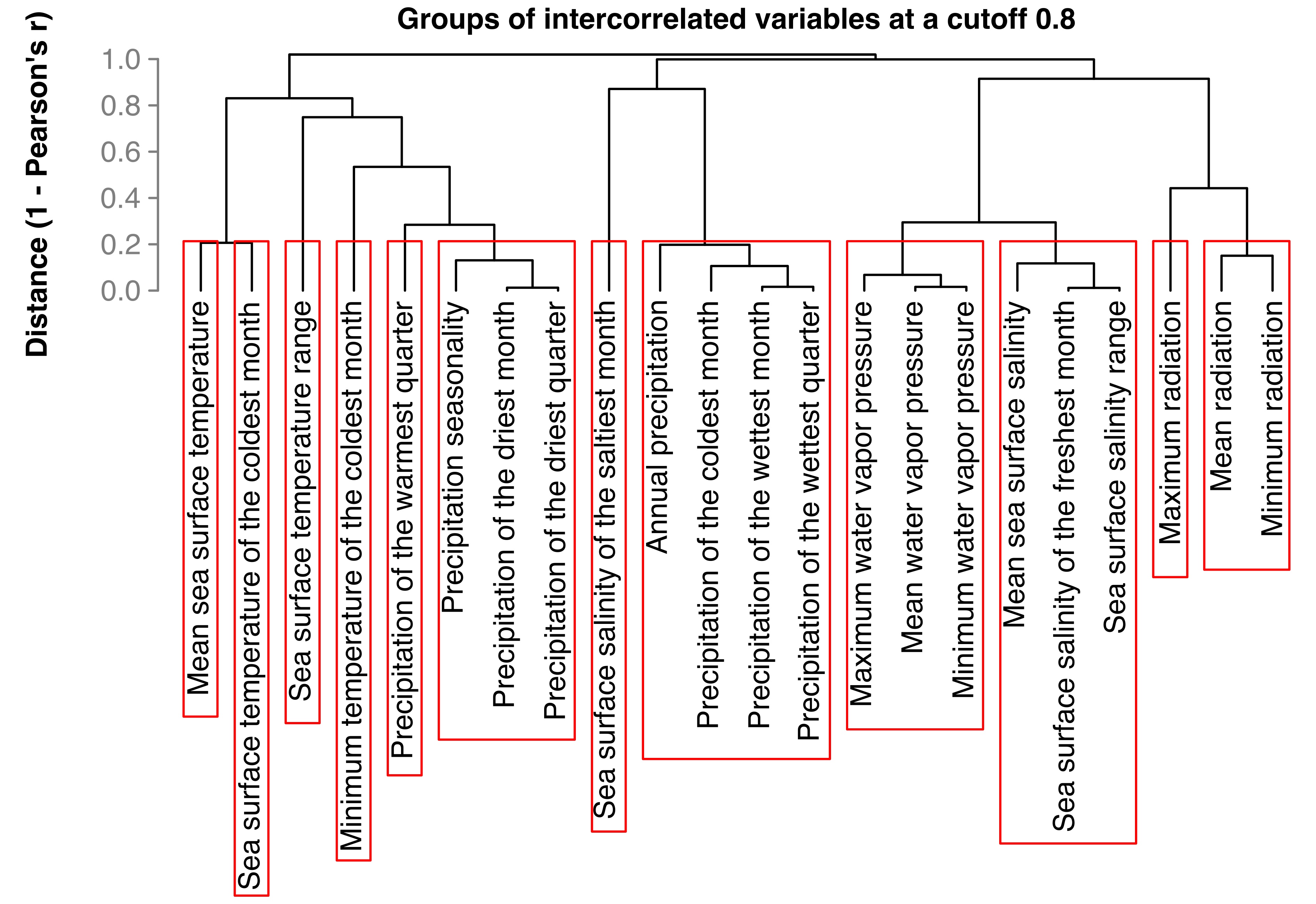

**Figure S1. Groups of collinear climatic and oceanographic variables from the WorldClim (Hijmans, Cameron, Parra, Jones, & Jarvis, 2005) and Marspec (Sbrocco & Barber, 2013) databases for genetic-environment association tests.** Pearson’s correlation cutoff was set to 0.8. Collinear variable groups are contained within red rectangles.

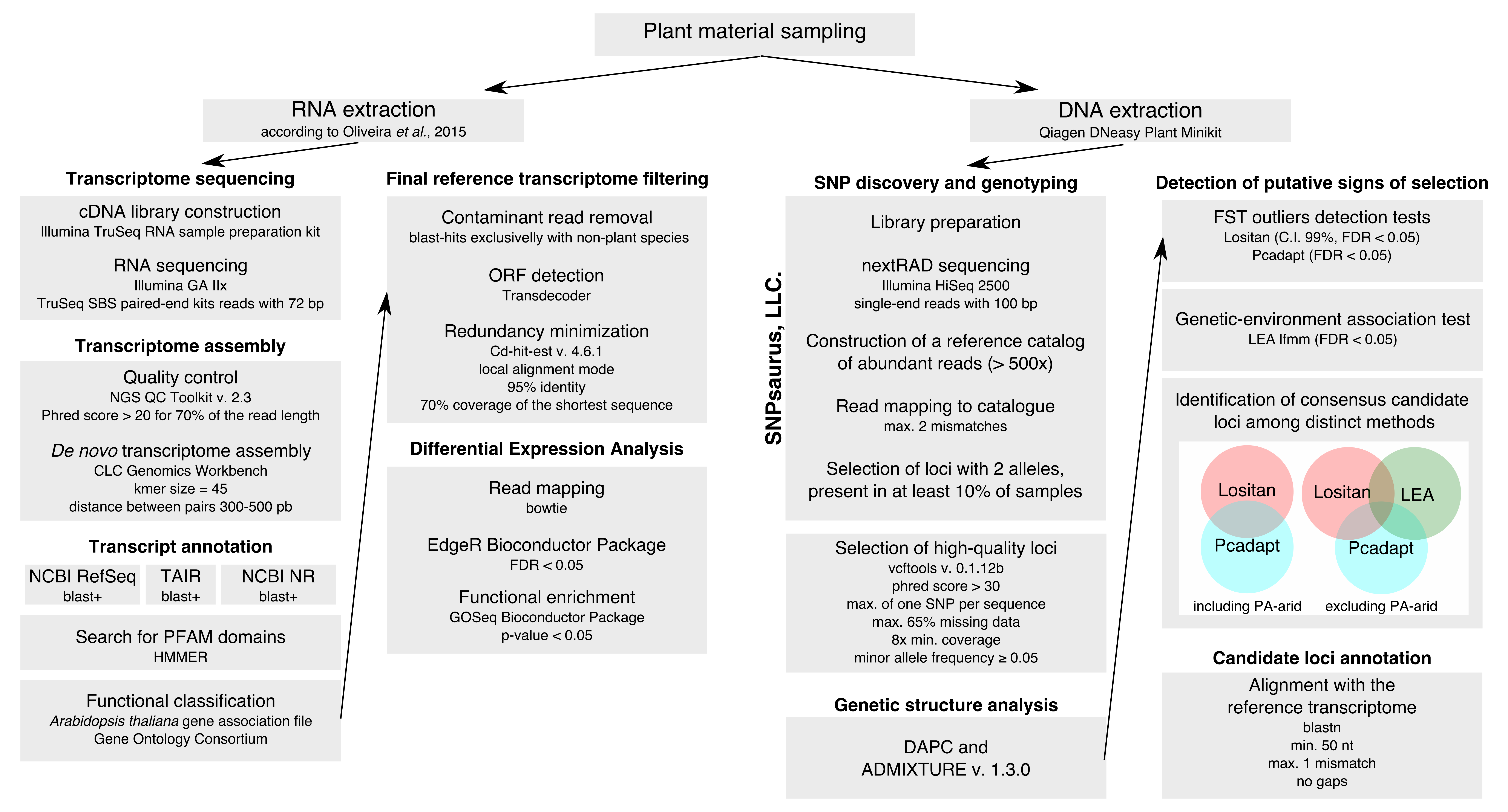

**Figure S2.** **Schematic representation of the methodological pipeline used in this study for RNA sequencing (RNA-Seq) and Nextera-tagmented reductively-amplified DNA sequencing (nextRAD).**

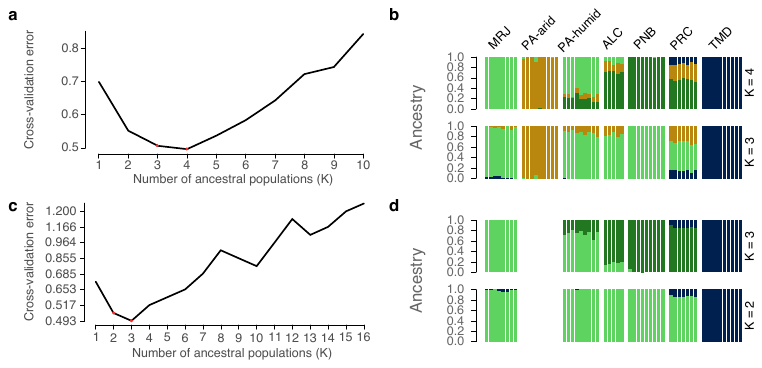

**Figure S3. Cross-validation error criteria for identifying the most likely number of ancestral populations and most likely scenarios in the program Admixture v. 1.3.0.** (a) Cross-validation plot for identifying the best K value from all sampled individuals. (b) Attribution of ancestry from the first (K = 4) and second (K = 3) most likely scenarios for all sampled individuals; stacked bars represent individuals, with each color representing one ancestral cluster. (c) Cross-validation plot for identifying the best K value from all sampled individuals, except for individuals sampled in the PA-arid site. (d) Attribution of ancestry form the first (K = 3) and second (K = 2) most likely scenarios for all sampled individuals, except for individuals sampled in the PA-arid site; stacked bars represent individuals, with each color representing one ancestral cluster.

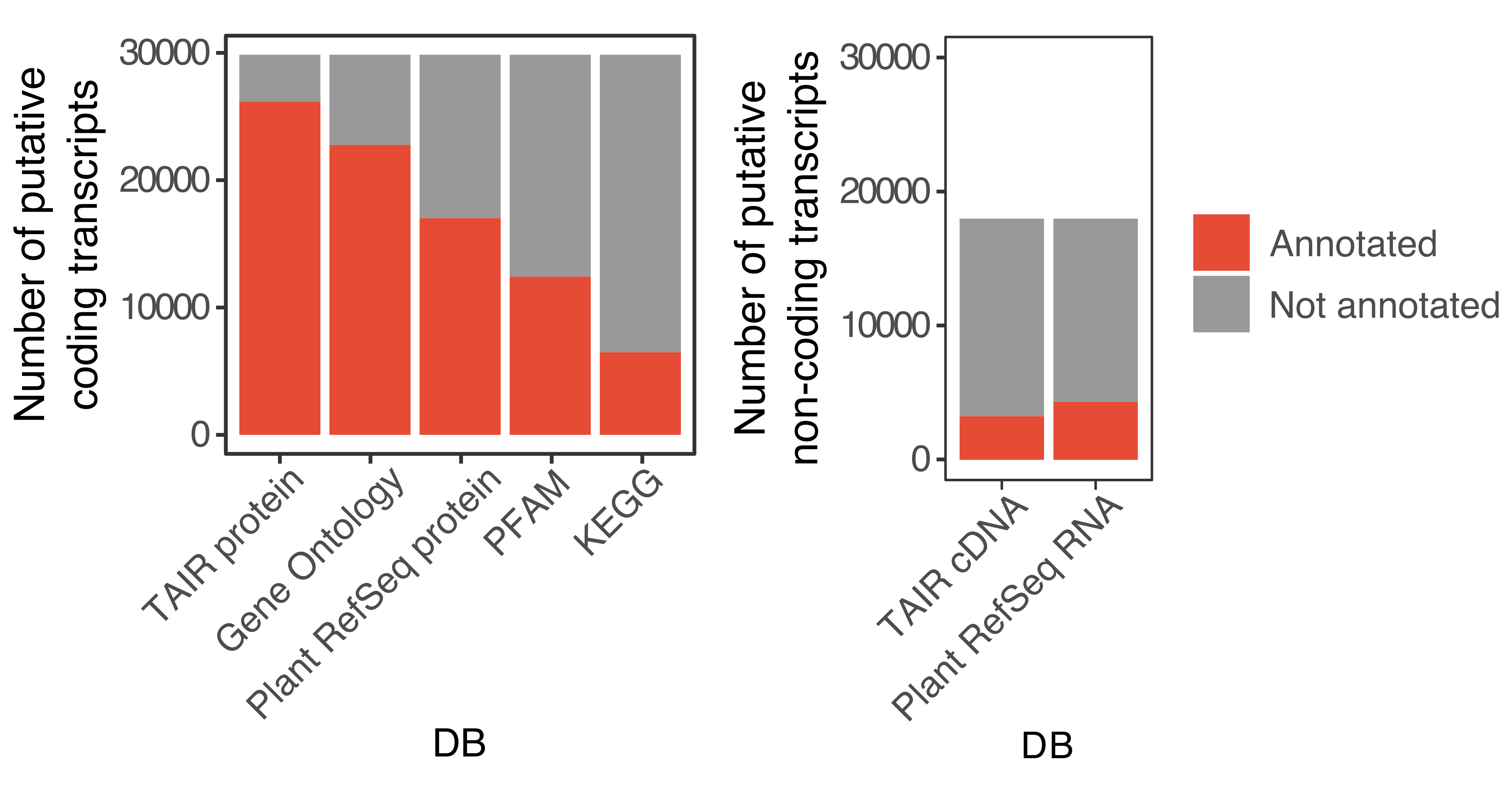

**Figure S4. Proportion of nonredundant transcripts of *Avicennia germinans* annotated using distinct reference databases (DB).** Red bars represent annotated transcripts, and gray bars represent nonannotated transcripts. *Left*: Annotation of transcripts containing an open reading frame (ORF) for protein synthesis (putative coding). *Right*: Annotation of putative non-coding transcripts.

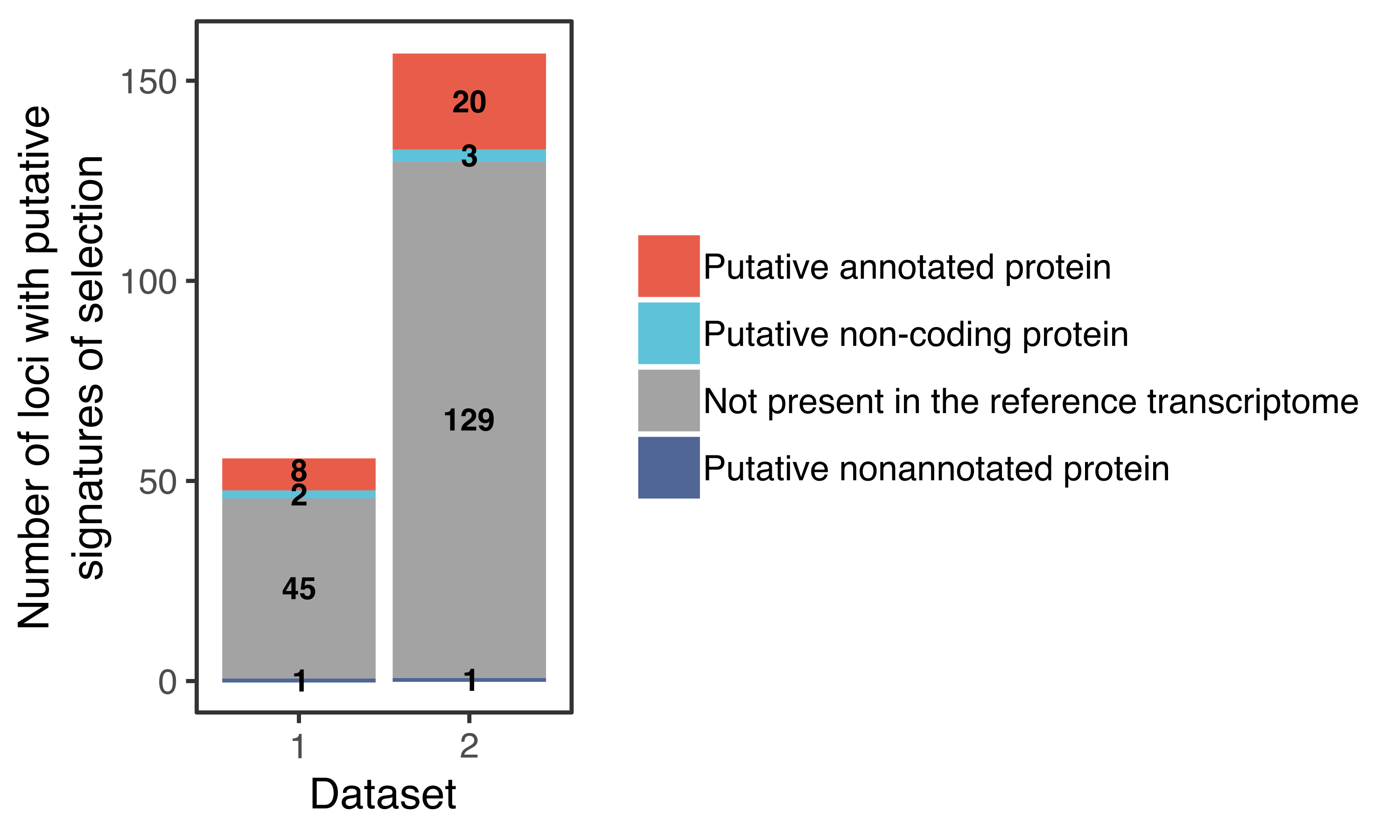

**Figure S5. Number of loci putatively under selection identified by at least two distinct methods.** Candidate loci were detected from two datasets: (1) all sampled individuals, including PA-arid samples, using F_ST_ outlier approaches and (2) a subset of samples, excluding individuals from the PA-arid site, combining genetic-environmental association tests and F_ST_ outlier tests.

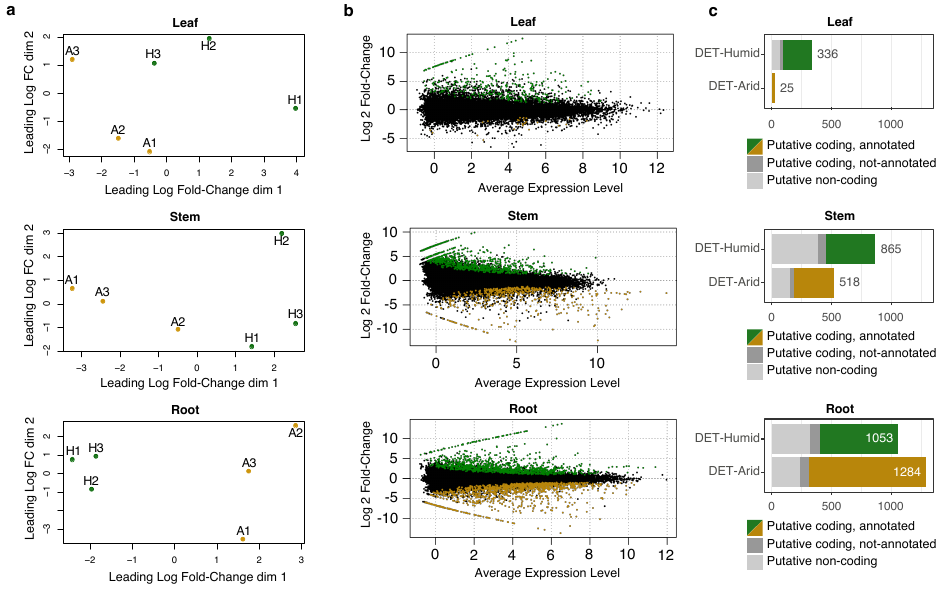

**Figure S6.** **Detection and annotation of differentially expressed transcripts (DETs).** (a) Two-dimensional scatterplot of distances among transcript expression profiles in leaf, stem and root samples (top to bottom) of *Avicennia germinans* seedlings acclimated in pots during three days in open-air, shaded and watered conditions; points represent individual samples, and distances are approximately the log2-transformed fold changes between samples; yellow points represent samples from the PA-arid site and green points represent samples from the PA-humid site. (b) MA-plots of transcript expression in leaves, stems and roots; yellow points represent DETs showing higher expression in samples from the PA-arid site than in samples from the PA-humid site (DET-Arid); green points represent DETs with higher expression in samples from the PA-humid than in samples from the PA-arid site (DET-Humid); black points represent non-differentially expressed transcripts. (c) Proportion of annotated and putative protein-coding DETs. Annotation of putative coding transcripts was performed using the blastx algorithm with *Arabidopsis thaliana* proteins as a reference; gray colored bars represent putative non-coding transcripts that are differentially expressed, and black bars represent putative coding transcripts that are differentially expressed but not annotated.

**2.	Tables**

**Table S1.** Pairwise population differentiation due to genetic structure, estimated by Wright’s fixation index (F_ST_, below diagonal gray cells), and by Nei’s standard genetic distance (above diagonal gray cells).

|  | MRJ | PAa | PAb | ALC | PNB | PRC | TMD |
| --- | --- | --- | --- | --- | --- | --- | --- |
| MRJ |  | 0.22 | 0.04 | 0.10 | 0.11 | 0.14 | 0.28 |
| PAa | 0.31 |  | 0.18 | 0.20 | 0.27 | 0.18 | 0.47 |
| PAb | 0.04 | 0.25 |  | 0.07 | 0.07 | 0.11 | 0.28 |
| ALC | 0.16 | 0.26 | 0.09 |  | 0.05 | 0.08 | 0.32 |
| PNB | 0.23 | 0.39 | 0.15 | 0.08 |  | 0.09 | 0.31 |
| PRC | 0.21 | 0.23 | 0.15 | 0.09 | 0.16 |  | 0.26 |
| TMD | 0.52 | 0.58 | 0.49 | 0.56 | 0.58 | 0.46 |  |

**Table S2.** Characterization of the *de novo-*assembled reference transcriptome of *Avicennia germinans*.

|  | All contigs | Putative coding contigs | Putative non-coding contigs |
| --- | --- | --- | --- |
| Number of contigs | 47,821 | 29,854 | 17,967 |
| Total size (bp) | 45,317,158 | 36,563,009 | 8,754,138 |
| Shortest contig (bp) | 294 | 298 | 294 |
| Longest contig (bp) | 43,423 | 43,423 | 5,755 |
| Number of contigs < 1Kb | 32,255 | 14,917 | 17,338 |
| Number of contigs ≥ 1Kb | 15,566 | 14,937 | 629 |
| Average contig size (bp) | 947.6 | 1,224.7 | 487.2 |
| Median contig size (bp) | 649 | 1,000 | 410 |
| Contig N50 (bp) | 1,327 | 1,588 | 482 |
| Average ORF size (amino acid) | NA | 307 | NA |
| Median ORF size (amino acid) | NA | 228 | NA |
| Longest ORF (amino acid) | NA | 3,753 | NA |
| Shortest ORF (amino acid) | NA | 100 | NA |

**Table S3**. Quality parameters of the *Avicennia germinans* transcriptome assembly.

| Reads mapping to contiguous sequences in the transcriptome | Mapped to a single contig | 82.40% |
| --- | --- | --- |
|  | Mapped to more than one contig | 1.31% |
|  | Not mapped to the transcriptome | 16.30% |
| BUSCO analysis of plant universal single-copy orthologs present in the transcriptome | Complete orthologous sequences present in single copy in the transcriptome | 63.49% |
|  | Complete orthologous sequences present in more than one copy in the transcriptome | 19.04% |
|  | Fragmented orthologous sequences present in the transcriptome | 9.21% |
|  | Absent orthologous sequences in the transcriptome | 8.26% |

**Table S4.** Putative orthologous sequences between *Avicennia germinans* transcriptome obtained from roots, stems and leaves and publicly available transcriptomes from other *Avicennia* L. (Avicenniaceae) species.

| Species | Plant material | NCBI accession number | Number of transcripts | Total number of putative orthologs |
| --- | --- | --- | --- | --- |
| *Avicennia marina* | leaves | GBIO01000000 | 89,833 | 29,496 |
| *Avicennia officinalis* | leaves | GFLY01000000 | 38,756 | 20,529 |
| *Avicennia officinalis* | roots | GSE73807 | 121,929 | 27,690 |
| *Avicennia schaueriana* | leaves, stems and flowers | GSE116060 | 49,490 | 31,348 |

**Table S5** is too large to be displayed in this file. It was submitted as a separate Supplemental information file.
